# MethylHDA: One-pot helicase-dependent amplification with concurrent DNMT1-mediated maintenance of DNA methylation

**DOI:** 10.64898/2026.03.05.709983

**Authors:** Kiat Whye Kong, Si En Poh, Fong Tian Wong, Yiqi Seow, Winston Lian Chye Koh

## Abstract

DNA methylation is an important epigenetic modification involved in gene regulation, genome stability, development, and disease. Analysis of low-abundance methylated DNA often requires DNA amplification. Conventional amplification methods, however, do not preserve methylation marks on newly synthesized DNA, potentially compromising downstream methylation analysis. Recently, DNA methyltransferase 1 (DNMT1) was coupled with phi29-mediated whole-genome amplification to preserve DNA methylation for downstream bisulfite sequencing. Targeted amplification of short, predefined genomic loci requires a different amplification strategy. Here, we developed MethylHDA, a one-pot isothermal workflow that integrates primer-directed helicase-dependent amplification (HDA) with DNMT1-mediated maintenance of DNA methylation. Because standard thermophilic HDA is performed at approximately 65 °C, a temperature incompatible with sustained DNMT1 activity, we established a unified reaction buffer and operating temperature of 42 °C that supported both HDA and DNMT1. Under these conditions, HDA produced an approximately five-cycle reduction in qPCR threshold cycle (Ct) relative to the unamplified control. Methylation-sensitive restriction enzyme quantitative PCR (MSRE-qPCR) demonstrated that inclusion of DNMT1 increased retention of methylation-dependent restriction protection and that the methylation signal was proportional to the methylated fraction of the input templates. These results demonstrate the feasibility of coupling targeted isothermal DNA amplification with concurrent maintenance of DNA methylation in a single reaction. This approach provides a foundation for methylation-preserving preamplification of short, locus-specific targets for downstream MSRE-qPCR and other targeted methylation assays.

## Introduction

DNA methylation at the C5 position of cytosine within CpG dinucleotides is a fundamental epigenetic modification that regulates gene expression and preserves genomic stability, contributing to transcriptional silencing, X-chromosome inactivation, and genomic imprinting (Jaenisch and Bird, 2003; Smith and Meissner, 2013; Moosavi and Ardekani, 2016; Jin and Liu, 2018; Michalak et al., 2019). Aberrant methylation patterns are closely associated with cancer, neurodegenerative disorders, and age-related diseases (Baylin and Jones, 2016; Moosavi and Ardekani, 2016; Jin and Liu, 2018; Michalak et al., 2019), underscoring the importance of accurate and sensitive methylation analysis.

DNA methylation can be analysed using genome-scale approaches, such as DNA methylation arrays and whole-genome sequencing-based methods, or targeted assays including methylation-specific PCR, pyrosequencing, methylation-sensitive high-resolution melting, methylation-specific multiplex ligation-dependent probe amplification, methylation-sensitive restriction enzyme quantitative PCR (MSRE-qPCR), and other locus-specific methylation assays (Waalwijk and Flavell, 1978; Singer et al., 1979; Herman et al., 1996; Eads et al., 2000; Colella et al., 2003; Nygren et al., 2005; Wojdacz and Dobrovic, 2007; Levenson, 2010; Holmes et al., 2014; Leontiou et al., 2015; Kint et al., 2018; Liu et al., 2019; Hong and Shin, 2021; Sun et al., 2021; Bendixen et al., 2023; Peters et al., 2024). Genome-scale approaches are valuable for methylome discovery but often require substantial DNA input, specialized instrumentation, and extensive downstream analysis (Holmes et al., 2014; Leontiou et al., 2015; Kint et al., 2018; Liu et al., 2019; Hong and Shin, 2021; Sun et al., 2021; Peters et al., 2024). By comparison, targeted assays are well suited for analysing predefined methylation biomarkers but can be limited by the amount of available DNA or the proportion of methylated target molecules within a sample (Herman et al., 1996; Eads et al., 2000; Colella et al., 2003; Nygren et al., 2005; Wojdacz and Dobrovic, 2007; Bendixen et al., 2023). This is particularly relevant for challenging specimens such as cell-free DNA, small biopsies, and other limited or fragmented clinical samples (Wan et al., 2017; Heitzer et al., 2019).

Preamplification could increase the amount of target DNA available for downstream methylation analysis, but conventional DNA amplification propagates nucleotide sequence without copying the associated cytosine methylation marks. During amplification of a methylated template, the parental strand retains its methylation while the newly synthesized strand is initially unmethylated, producing a hemi-methylated intermediate (Pradhan et al., 1999; Goyal et al., 2006; Smith and Meissner, 2013; Liu et al., 2020). In the absence of a maintenance methyltransferase, subsequent rounds of DNA synthesis progressively dilute the original methylation signal among unmethylated amplification products (Goyal et al., 2006; Liu et al., 2020). Consequently, the methylated fraction measured after amplification may no longer accurately represent that of the input sample.

DNA methyltransferase 1 (DNMT1) preferentially methylates hemi-methylated CpG sites and is the principal enzyme responsible for maintenance of DNA methylation following DNA replication in mammalian cells (Pradhan et al., 1999; Goyal et al., 2006; Pradhan et al., 2008; Smith and Meissner, 2013; Schrader et al., 2015; Adam et al., 2020; Liu et al., 2020; New England Biolabs, n.d.). Liu *et al*. exploited this activity in a methylation-preserving whole-genome amplification method, termed 5mC-WGA, which combined DNMT1 with phi29 DNA polymerase-mediated multiple-displacement amplification (Liu et al., 2020). Their study demonstrated that DNA synthesis and maintenance of DNA methylation could be coupled *in vitro* and enabled whole-genome methylome analysis from limited DNA inputs using downstream bisulfite sequencing (Liu et al., 2020). Phi29-mediated multiple-displacement amplification is well suited for whole-genome amplification because it combines random priming, strong strand-displacement activity, and highly processive DNA synthesis (Liu et al., 2020). In contrast, targeted methylation assays require selective amplification of short, predefined genomic regions containing clinically or biologically relevant methylation biomarkers or methylation-sensitive restriction sites. A DNMT1-coupled amplification strategy designed specifically for targeted methylation analysis has not yet been established.

Helicase-dependent amplification (HDA) is a sequence-directed isothermal amplification method in which a DNA helicase separates duplex DNA to permit primer annealing and polymerase-mediated strand synthesis (Vincent et al., 2004; Gill and Ghaemi, 2008; Jeong et al., 2009). Unlike random-primer whole-genome amplification, the boundaries of an HDA product are defined by a pair of sequence-specific primers (Vincent et al., 2004; Gill and Ghaemi, 2008; Jeong et al., 2009). HDA therefore enables selective amplification of short genomic regions surrounding predefined CpG sites while avoiding thermocycling and the generation of complex whole-genome amplification products. These characteristics make HDA an attractive preamplification strategy for targeted methylation assays such as MSRE-qPCR. The use of short, primer-defined amplicons may also facilitate analysis of fragmented DNA (Wan et al., 2017; Heitzer et al., 2019) and the development of compact locus-specific methylation panels, although these applications require further validation.

Coupling HDA with DNMT1 nevertheless presents a substantial biochemical compatibility challenge. In this study, HDA was implemented using the commercially available IsoAmp® II Universal thermophilic HDA system, which is recommended to operate at 65 °C (New England Biolabs, n.d.). This temperature is incompatible with sustained activity of mammalian DNMT1, which normally functions near physiological temperatures and progressively loses activity at elevated temperatures (Pradhan et al., 1999; Pradhan et al., 2008; Syeda et al., 2011; Schrader et al., 2015; BioTreks, 2016; Adam et al., 2020; New England Biolabs, n.d.). Although engineering a thermostable DNMT1 could potentially overcome this incompatibility (iGEM Team Heidelberg, 2014), we instead sought to identify a common reaction condition that would support both enzymes without protein engineering.

Here, we established a unified reaction buffer and reduced-temperature HDA conditions that supported both targeted DNA amplification and DNMT1-mediated maintenance of DNA methylation within a single isothermal reaction. Reaction conditions were optimized to balance efficient DNA amplification, preservation of methylation in products derived from methylated templates, and suppression of DNMT1-mediated methylation of products derived from unmethylated templates. The resulting one-pot workflow was evaluated using MSRE-qPCR to determine whether input-dependent methylation signals could be retained following amplification. This study establishes proof of concept for a targeted HDA–DNMT1 workflow and provides a foundation for methylation-preserving preamplification of predefined genomic loci for downstream MSRE-qPCR and other targeted methylation assays.

## Results

### Concept and design of concurrent DNA amplification and DNMT1-mediated methylation

Establishing a targeted methylation-preserving amplification workflow requires integration of helicase-dependent amplification (HDA) with DNA methyltransferase 1 (DNMT1)-mediated maintenance of DNA methylation in a single isothermal reaction. Successful implementation depends on reaction conditions that simultaneously support DNA amplification and DNMT1 activity while preserving the methylation status of the input template. The proposed workflow relies on the generation of hemi-methylated amplification intermediates during HDA, which serve as substrates for DNMT1-mediated maintenance methylation (Figure 1a).

**Figure 1.**
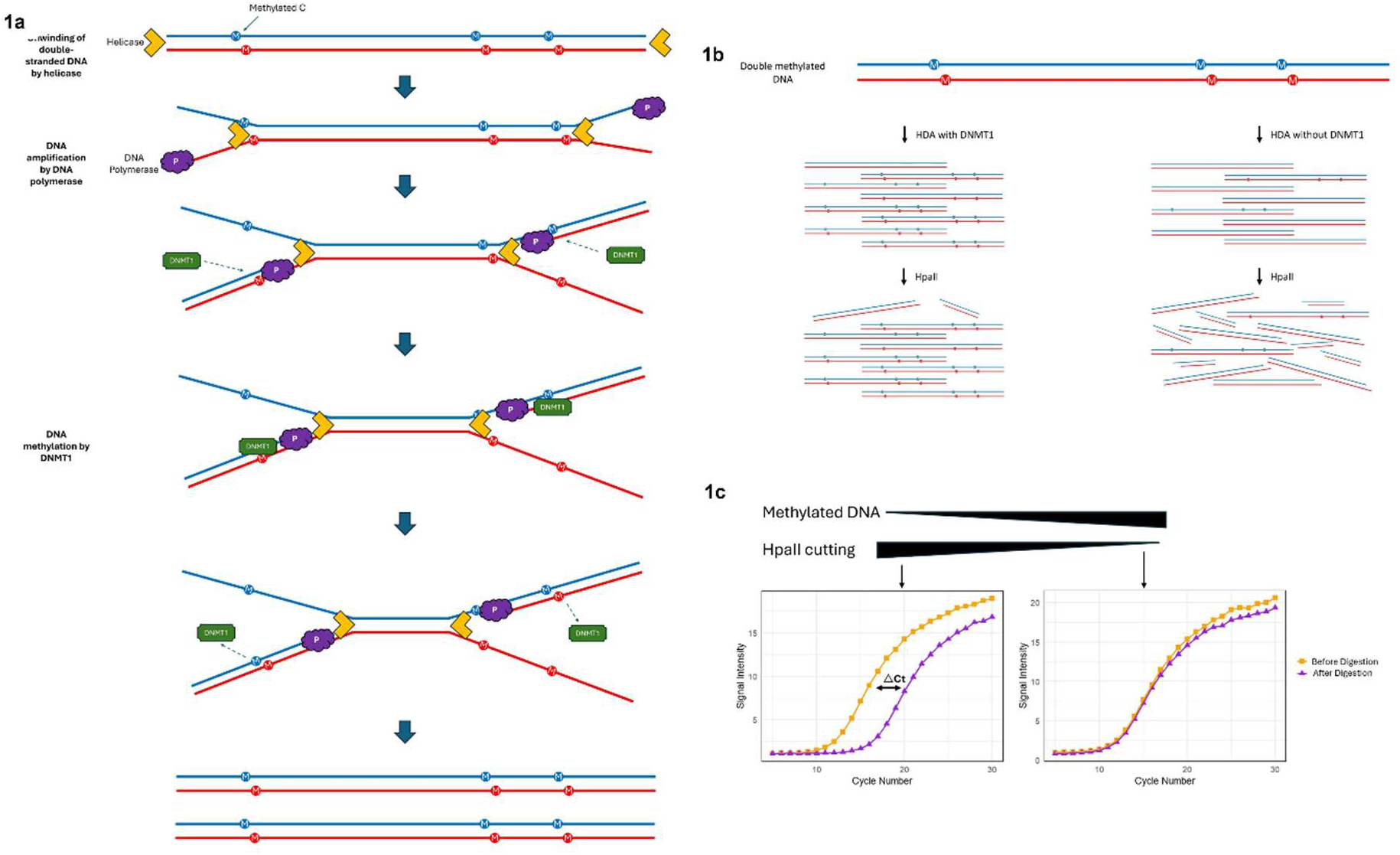
Schematic model and assay for one-pot DNA amplification and DNMT1-mediated methylation. (a) Proposed mechanism illustrating concurrent DNA amplification and methylation. Double-stranded methylated DNA undergoes a one-pot reaction containing helicase, DNA polymerase, and DNMT1. Helicase unwinds the duplex, exposing single-stranded templates for polymerase-driven synthesis. DNMT1 recognizes hemi-methylated CpG sites and methylates the nascent strand, yielding double-stranded products expected to undergo DNMT1-mediated maintenance of DNA methylation. (b) Workflow of the one-pot amplification–methylation assay and downstream analysis. HDA is performed in the presence or absence of DNMT1. In reactions containing DNMT1, newly synthesized strands are methylated at CpG sites, protecting the products from HpaII digestion. Without DNMT1, amplification occurs, but the DNA remains unmethylated and is cleaved by HpaII. (c) Expected MSRE-qPCR outcomes following HpaII digestion. Methylated products resist cleavage and show low ΔCt values, whereas unmethylated products are digested, resulting in delayed qPCR amplification and higher ΔCt values.

A major challenge is the different operating requirements of the two enzymatic systems. Mammalian DNMT1 functions most efficiently near physiological temperatures and is progressively inactivated at the elevated temperatures used for conventional PCR (Pradhan et al., 1999; Pradhan et al., 2008; Schrader et al., 2015; BioTreks, 2016; Adam et al., 2020; New England Biolabs, n.d.). We therefore employed the commercially available IsoAmp® II Universal thermophilic HDA (tHDA) system, which performs DNA amplification under constant-temperature conditions and provides a suitable platform for identifying reaction conditions compatible with both HDA and DNMT1. This strategy enables helicase-mediated strand separation, primer-directed DNA synthesis, and DNMT1-mediated maintenance methylation to occur within a single reaction.

As illustrated in Figure 1a, helicase unwinds the DNA duplex to expose single-stranded regions, which are copied by DNA polymerase. Simultaneously, DNMT1 targets hemi-methylated CpG sites generated during synthesis and methylates the complementary strand, effectively replicating both the sequence and methylation status of the original template. Amplification in the presence of DNMT1 is expected to produce newly synthesized strands methylated at CpG sites corresponding to the template, protecting them from cleavage by HpaII, a methylation-sensitive restriction enzyme (MSRE) that selectively digests unmethylated CCGG sequences (Figure 1b). The methylation status of the amplified DNA can be quantified using MSRE-quantitative PCR (qPCR), where the difference in Ct values before and after HpaII digestion (ΔCt) reflects the extent of methylation (Figure 1c). A small ΔCt indicates protection from digestion due to DNMT1-mediated methylation, whereas a large ΔCt corresponds to extensive cleavage of unmethylated DNA. This integrated isothermal reaction is therefore designed to generate double-stranded DNA products that maintain DNA methylation during amplification based on the methylation status of the input template, while reducing sample handling and eliminating the need for separate processing steps.

### Assessment of DNMT1 catalytic activity above its optimum temperature for compatibility with isothermal amplification

DNMT1 exhibits optimal catalytic activity at 37 °C but loses stability at temperatures above 60 °C (Pradhan et al., 1999; Pradhan et al., 2008; Schrader et al., 2015; BioTreks, 2016; Adam et al., 2020; New England Biolabs, n.d.). For the amplification component, we used the IsoAmp® Enzyme Mix from the IsoAmp® II Universal tHDA Kit. Although the IsoAmp® Enzyme Mix has a recommended operating temperature of 65 °C, we aimed to determine whether DNMT1 retains sufficient activity at temperatures above its optimal 37 °C that could potentially overlap with the working range of the IsoAmp® enzymes, enabling a one-pot amplification and methylation reaction.

To evaluate DNMT1 activity across temperatures, we adapted a qPCR-based assay involving methylation-dependent restriction enzyme digestion, originally described by Petell et al. (2017). This assay is sensitive to CpG methylation and allows quantification of methylation efficiency under different conditions. In this study, we modified the DNA template so that the forward strand lacked an MspJI recognition site, while the reverse strand contained a site that becomes recognizable by MspJI only upon methylation, effectively creating a hemi-methylated duplex (Table 1). This design allowed us to specifically monitor DNMT1 activity because methylation of the reverse strand converts the duplex into a fully methylated substrate that can be cleaved by MspJI. Cleavage reduces the amount of amplifiable DNA, producing a higher Ct value in subsequent qPCR.

**Table 1.**
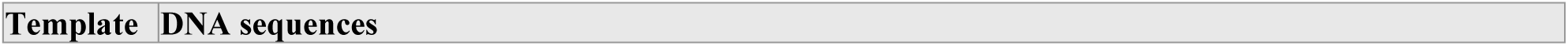
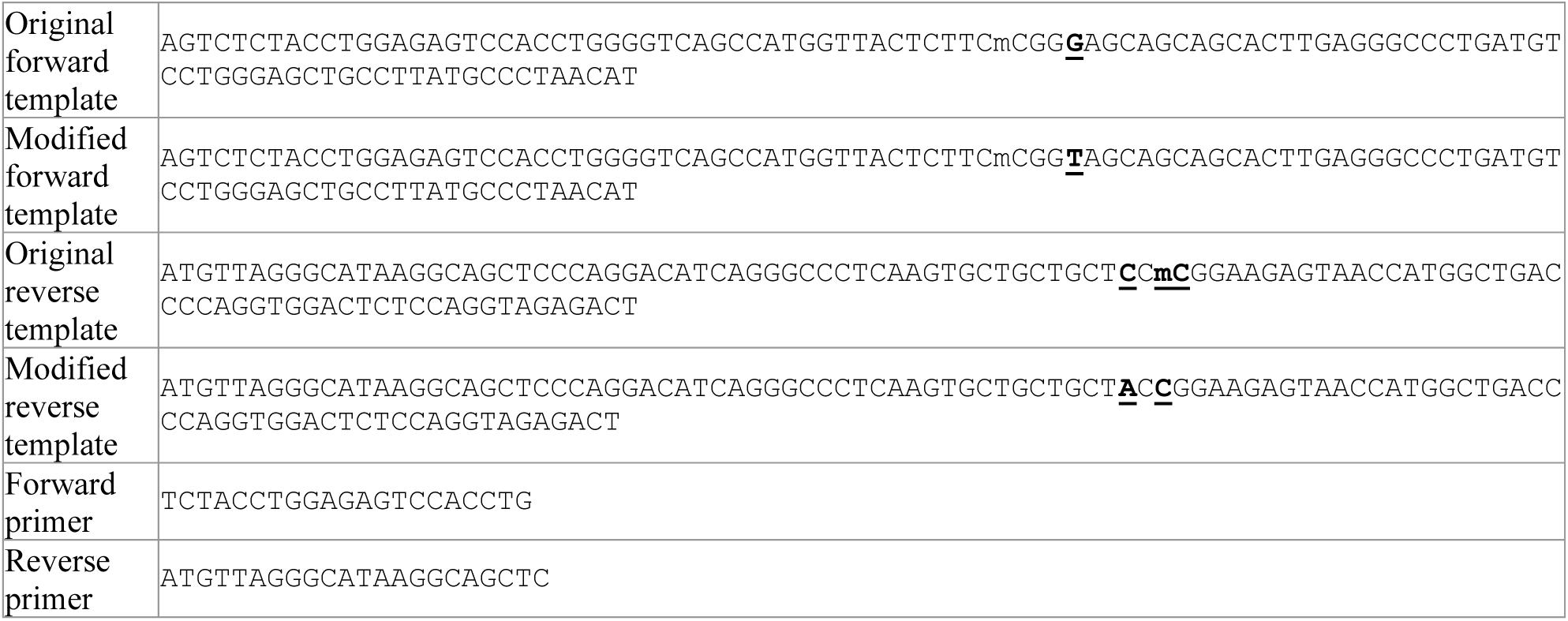
Template and primer sequences adapted from Petell et al. (2017) and modified template sequences (GenBank accession number PZ577457).

Using this system, DNMT1 reactions were carried out at 40 °C, 42 °C, and 45 °C. As shown in Figure 2a, all DNMT1-treated samples displayed a rightward shift in Ct following MspJI digestion, indicating successful methylation and subsequent cleavage. In contrast, water-treated controls lacking DNMT1 showed minimal Ct changes, confirming that the digestion was dependent on methylation. Comparison of the Ct shifts revealed that DNMT1 activity peaked at 42 °C, with lower efficiency observed at 40 °C and 45 °C. These results demonstrate that DNMT1 retains sufficient catalytic function at 42 °C, suggesting that this temperature could potentially be compatible with HDA.

**Figure 2.**
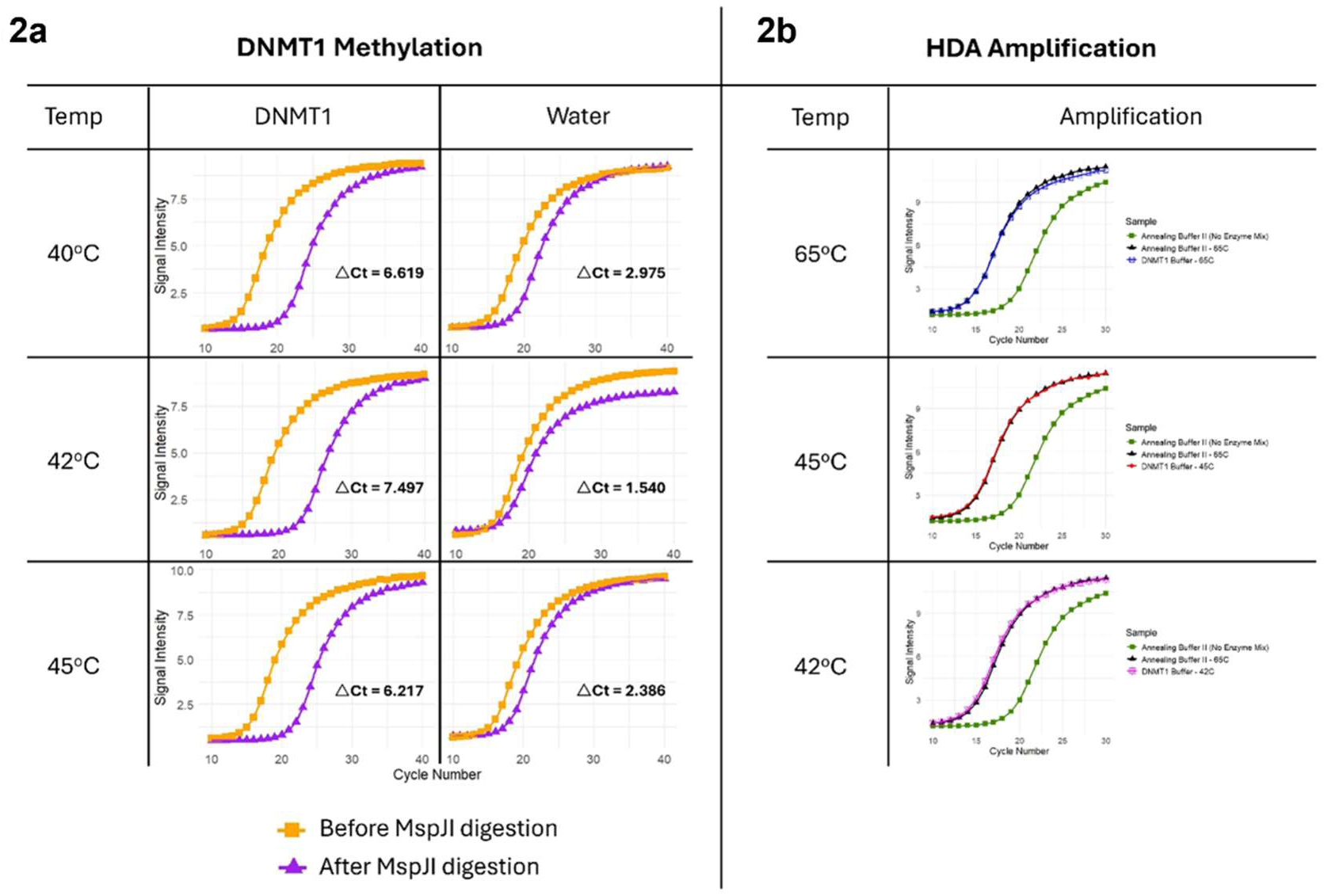
Temperature compatibility of DNMT1 methylation activity and IsoAmp® Enzyme Mix amplification efficiency under DNMT1-buffer conditions. (a) qPCR amplification plots of MspJI-digested DNA following DNMT1-mediated methylation at 40 °C, 42 °C, and 45 °C. Hemi-methylated DNA templates were incubated with DNMT1, then digested with MspJI. The forward strand was fully methylated, while the reverse strand was unmethylated and contained MspJI recognition sites. Successful methylation of the reverse strand by DNMT1 enables MspJI cleavage, resulting in decreased template availability and increased Ct values. Water-only controls (without DNMT1) served as negative controls, confirmed that digestion was methylation-dependent. The greatest rightward Ct shift relative to controls was observed at 42 °C, indicating peak DNMT1 activity. (b) Amplification efficiency of the IsoAmp® Enzyme Mix in DNMT1 buffer compared with standard Annealing Buffer II at 65 °C, 45 °C, and 42 °C.

Reactions included positive controls (Annealing Buffer II at 65 °C with IsoAmp® Enzyme Mix), negative controls (without IsoAmp® Enzyme Mix), and DNMT1-buffer reactions (with IsoAmp® Enzyme Mix supplemented with BSA and SAM) using 100% double-methylated template. Comparable amplification between DNMT1 buffer and Annealing Buffer II at 65 °C demonstrates that the IsoAmp® Enzyme Mix performs efficiently under DNMT1-compatible conditions. Amplification at 42 °C and 45 °C in DNMT1 buffer was similar to the positive control, confirming robust enzyme activity at temperatures suitable for DNMT1.

### Establishing HDA activity in DNMT1-compatible buffer and temperature ranges

Earlier, we established that DNMT1 retains catalytic activity at 42 °C, and we next sought to determine whether HDA, using IsoAmp® Enzyme Mix, could function effectively at these lower temperatures. Since our goal is to adapt HDA to operate under conditions compatible with DNMT1, it was essential to verify that the amplification system remains active in DNMT1-compatible buffer and at temperatures below its recommended temperature of 65 °C.

To assess compatibility with DNMT1-compatible conditions, HDA was performed using a modified double-methylated DNA template (from the MethylMiner™ Methylated DNA Enrichment Kit; Table 2) in DNMT1 buffer supplemented with Bovine Serum Albumin (BSA) and S-adenosyl methionine (SAM) across a temperature range of 42 °C to 65 °C. Amplification efficiency was compared to the standard IsoAmp® II Universal tHDA Kit protocol using Annealing Buffer II at 65 °C. Comparable amplification under both conditions would indicate that the HDA enzymes function effectively in DNMT1 buffer.

**Table 2.**
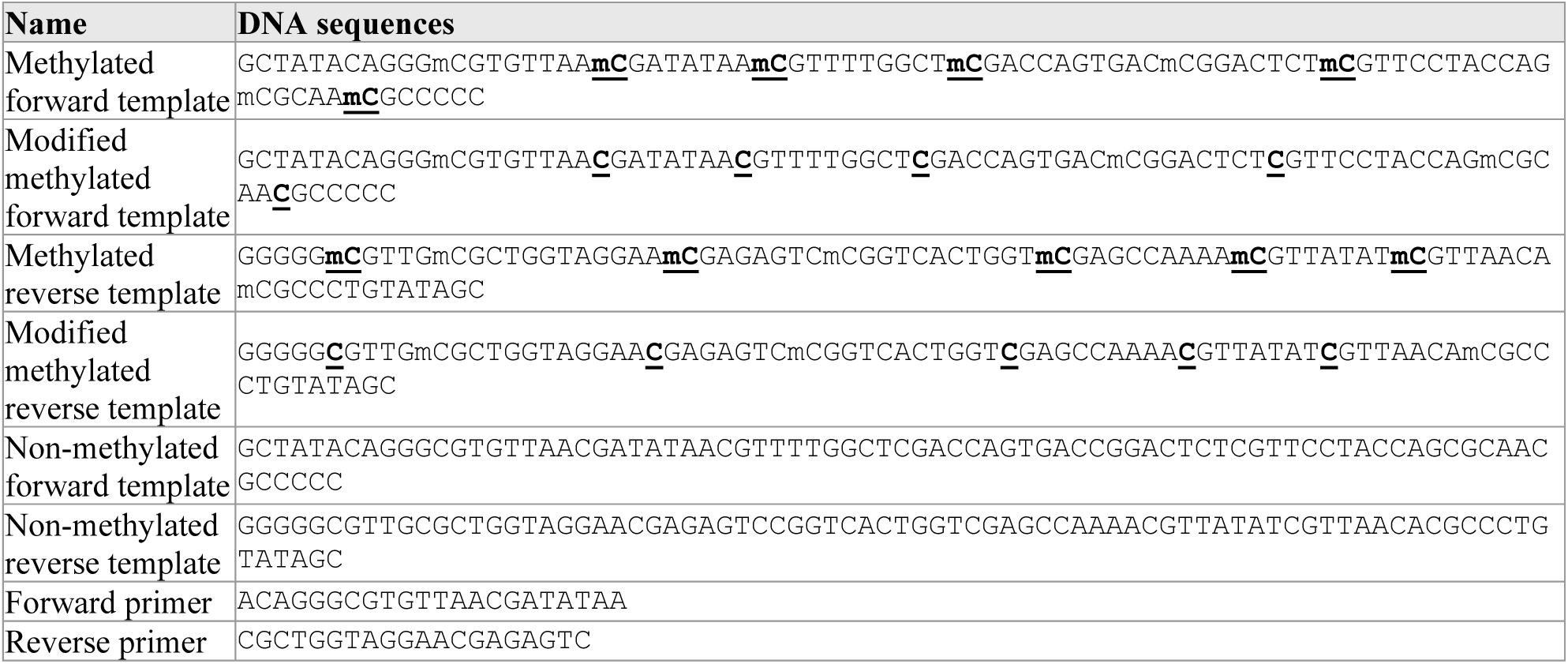
Template and primer sequences adapted from MethylMiner™ Methylated DNA Enrichment Kit and modified template sequences (GenBank accession number PZ577458).

As shown in Figure 2b, amplification in Annealing Buffer II at 65 °C produced a clear Ct shift relative to the no-enzyme control, confirming robust DNA amplification. Amplification in DNMT1 buffer at the same temperature was similarly efficient, demonstrating that the HDA enzymes are compatible with DNMT1 reaction conditions. To evaluate amplification at temperatures more favourable for DNMT1, reactions were performed at 42 °C and 45 °C in DNMT1 buffer. Amplification at these temperatures was nearly equivalent to that in Annealing Buffer II at 65 °C. Together, these results indicate that HDA using the IsoAmp® Enzyme Mix remains active at 42 °C under DNMT1-compatible conditions.

### HDA proceeds efficiently under overlapping conditions in the final one-pot assay at 42 °C

Given that DNMT1 methylation was most effective at 42 °C, and that HDA could be performed at this temperature under DNMT1-compatible conditions with comparable efficiency to the manufacturer’s protocol, we proceeded to validate the amplification component of the one-pot assay. Template mixtures containing different ratios of modified methylated and unmethylated DNA from the MethylMiner™ Methylated DNA Enrichment Kit (Table 2) were subjected to amplification in the presence or absence of DNMT1 and the IsoAmp® Enzyme Mix. Amplification was monitored via qPCR, with reduced Ct values indicating successful DNA synthesis.

As shown in Figure 3, reactions containing the IsoAmp® Enzyme Mix (R1, R2, and R3) exhibited markedly lower Ct values, indicating efficient DNA amplification mediated by helicase-driven strand separation and polymerase activity. In contrast, reactions lacking the IsoAmp® Enzyme Mix (R4 and R5) produced higher Ct values, consistent with the absence of amplification. These results demonstrate that HDA proceeds efficiently under DNMT1-compatible buffer conditions at 42 °C, confirming that the amplification component functions as intended in the one-pot reaction.

**Figure 3.**
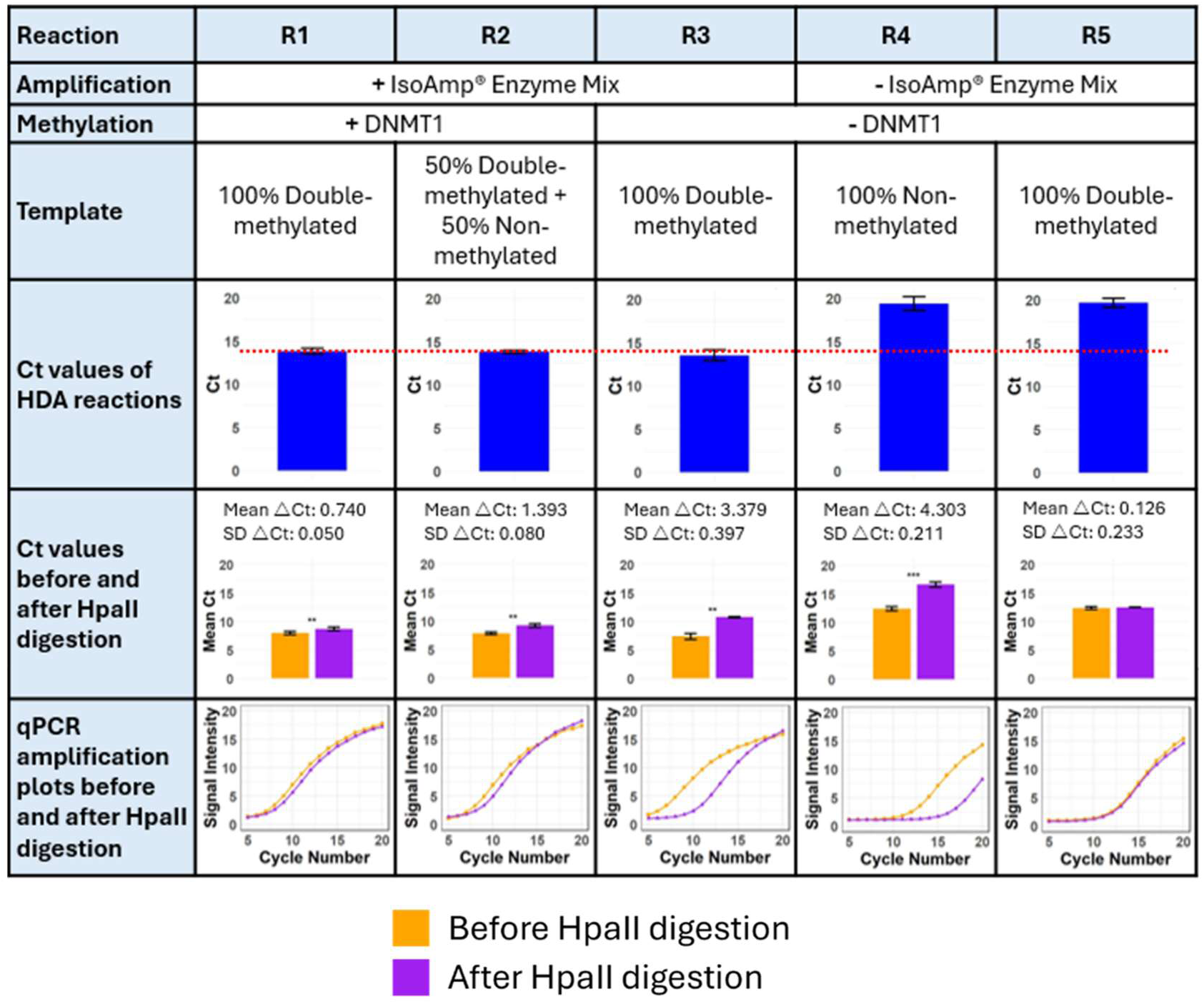
One-pot amplification and DNMT1-mediated methylation under DNMT1-compatible conditions at 42 °C. The table summarizes the combinations of DNA template with or without IsoAmp® Enzyme Mix and DNMT1 for each reaction, along with the corresponding qPCR analysis of HDA and DNMT1-mediated methylation. In reactions containing IsoAmp® Enzyme Mix (R1–R3), helicase-dependent amplification generated multiple DNA copies, resulting in lower Ct values. In contrast, reactions lacking IsoAmp® Enzyme Mix (R4 and R5) showed no amplification, leaving DNA templates amount unaltered and producing higher Ct values. A red dotted line on the bar chart indicates the threshold of successful amplification compared to no amplification. Bar charts and qPCR amplification plots before and after HpaII digestion show the results of DNMT1-mediated methylation during amplification. In R1, which contains DNMT1, the amplified product is methylated, protecting it from HpaII cleavage and resulting in minimal digestion, similar to the fully methylated DNA control (R5). In contrast, R3, lacking DNMT1, remains unmethylated and is digested like the non-methylated control (R4). R2, containing a 1:1 mix of double-methylated and non-methylated DNA, shows an intermediate ΔCt value, reflecting methylation levels proportional to the composition of the input template. These results demonstrate that DNMT1-mediated methylation occurs during HDA, with the extent of methylation corresponding to the methylation status of the starting templates as assessed by MSRE-qPCR.

### DNMT1-mediated methylation during one-pot HDA is proportional to the methylation status of the input template

After establishing efficient HDA under one-pot amplification–methylation conditions, we next evaluated DNMT1-mediated methylation within the same workflow using MSRE-qPCR. This was assessed by testing the susceptibility of the amplified products to digestion by the MSRE HpaII. HpaII specifically cleaves unmethylated CCGG sequences, whereas methylated cytosines at these sites prevent enzymatic cleavage. The modified templates from the MethylMiner™ Methylated DNA Enrichment Kit (Table 2) contain a HpaII recognition site, such that methylation confers resistance to cleavage while unmethylated DNA is digested. Therefore, successful DNMT1-mediated methylation during amplification would result in minimal change in Ct values before and after HpaII digestion. In contrast, unmethylated amplified products would be cleaved, resulting in higher Ct values due to reduced template availability. To verify HpaII digestion efficiency, control reactions with fully methylated and unmethylated DNA at three concentrations (100 nM, 10 nM, and 1 nM) were included (see Supplementary Figure S1).

As shown in Figure 3 (and Supplementary Figure S2), reaction R1, amplified from a fully methylated template in the presence of DNMT1, exhibited minimal Ct change following HpaII digestion, indicating effective methylation and protection of the newly synthesized DNA. This protection mirrored that of R5, the positive control containing fully methylated DNA without amplification or additional methylation, suggesting that most of the amplified products in R1 were successfully methylated by DNMT1. In contrast, R3, which was also amplified from a fully methylated template but lacked DNMT1, displayed a substantial Ct shift post-digestion, similar to R4, the fully non-methylated control. This indicates that in the absence of DNMT1, the amplified DNA remains largely unmethylated and thus susceptible to HpaII cleavage. R2 was prepared using a 1:1 mixture of methylated and non-methylated templates and therefore contains only half the amount of methylatable substrate for DNMT1, based on the reaction scheme in Figure 1. We predicted that only half of the amplified products would be methylated, resulting in a Ct shift intermediate between R1 and R3. Consistent with this, R2 exhibited a moderate Ct shift in the presence of DNMT1, indicating partial protection from digestion and confirming that methylation levels were proportional to the input template composition.

Together, these results demonstrate that DNMT1-mediated methylation occurs during HDA under the optimized reaction conditions. Furthermore, the MSRE-qPCR results indicate that the extent of methylation is proportional to the methylation status of the input template, supporting the feasibility of integrating HDA with DNMT1-mediated methylation maintenance in a one-pot workflow.

## Discussion

This study demonstrates the feasibility of coupling primer-directed helicase-dependent amplification (HDA) with DNA methyltransferase 1 (DNMT1)-mediated maintenance of DNA methylation under a unified isothermal condition. By lowering the operating temperature of a thermophilic HDA system and identifying a shared buffer environment, we established a one-pot reaction in which both target amplification and DNMT1 activity were retained. The resulting MethylHDA workflow establishes proof of concept for integrating HDA with DNMT1 in a single isothermal reaction. As a targeted counterpart to the previously reported phi29-based methylation-preserving whole-genome amplification strategy (Liu et al., 2020), it provides a foundation for future development of methylation-sensitive restriction enzyme quantitative PCR (MSRE-qPCR)-based assays.

The feasibility of this approach relies on the formation of hemi-methylated amplification intermediates that serve as substrates for DNMT1 (Figure 1). During synthesis from a methylated template, the parental strand retains its original methylation whereas the newly synthesized strand is initially unmethylated. DNMT1 preferentially recognizes these hemi-methylated CpG sites and methylates the complementary daughter strand (Pradhan et al., 1999; Goyal et al., 2006; Adam et al., 2020), thereby maintaining DNA methylation during amplification. Consistent with the proposed mechanism, HpaII resistance reflected the methylation status of the input templates. Products generated in the presence of DNMT1 exhibited greater resistance to HpaII digestion than those generated in its absence, while reactions containing a 1:1 mixture of methylated and unmethylated templates produced an intermediate MSRE-qPCR signal between the fully methylated and fully unmethylated controls (Figure 3). Together, these findings suggest that the workflow can preserve information related to the methylated fraction of the input DNA under the conditions examined. Nevertheless, these results do not yet establish complete or base-resolution maintenance of DNA methylation across the entire amplicon, nor do they define the quantitative relationship between input methylation levels and the resulting amplification products across a broad range of methylation fractions.

A major challenge in developing MethylHDA was reconciling the different operating conditions of the amplification and methylation enzymes. The IsoAmp® II Universal tHDA system is optimized for amplification at approximately 65 °C (New England Biolabs, n.d.), whereas mammalian DNMT1 is typically assayed near 37 °C and progressively loses activity at elevated temperatures (Pradhan et al., 1999; Pradhan et al., 2008; Adam et al., 2020). We found that the functional ranges of the two systems overlapped sufficiently to support both activities at 42 °C (Figure 2). This finding demonstrates that enzyme optima do not necessarily define absolute operating limits and that a shared biochemical window can be established through joint optimization of temperature, buffer composition and reaction time. Nevertheless, 42 °C represents a compromise rather than the optimum for either enzyme, and the kinetics of DNA amplification and maintenance methylation under these conditions remain to be investigated.

MethylHDA differs from the previously described 5mC-WGA method (Liu et al., 2020) primarily in analytical scale and amplification architecture. The phi29-based method uses random priming and high-processivity strand-displacement synthesis to produce broad genomic amplification suitable for whole-methylome sequencing (Liu et al., 2020). In contrast, HDA uses sequence-specific primers to generate short products with defined boundaries. Accordingly, MethylHDA is intended for targeted analysis of preselected loci rather than unbiased whole-methylome coverage. These approaches should be viewed as complementary rather than competing: phi29–DNMT1 amplification is appropriate when genome-scale methylation information is required, whereas HDA–DNMT1 may be advantageous when the analytical objective is to enrich one or a small number of known methylation biomarkers prior to targeted analysis.

The use of short, primer-defined amplicons provides several potential advantages. It confines amplification to the region containing the methylation-sensitive site, reduces the complexity of the amplification products and facilitates coupling with MSRE-qPCR. Short amplicons also reduce dependence on long, intact DNA templates and may therefore prove advantageous for fragmented specimens such as circulating cell-free DNA or formalin-fixed material (Wan et al., 2017; Heitzer et al., 2019). These applications were not evaluated in the present study, which used synthetic DNA templates, and will require validation using biologically and clinically relevant samples. Furthermore, because HDA does not require repeated high-temperature denaturation, it is inherently more compatible with mammalian DNMT1 than conventional PCR.

The MethylHDA reaction concept may also be extended to other isothermal amplification platforms. Recombinase polymerase amplification (RPA) (Tan et al., 2022) operates within a temperature range compatible with mammalian DNMT1, making it an attractive candidate for future methylation-preserving amplification workflows. However, its recombinase, single-stranded DNA-binding proteins, strand-displacing polymerase and reaction environment may influence DNMT1 accessibility to hemi-methylated substrates and therefore require experimental evaluation. In contrast, loop-mediated isothermal amplification (LAMP) (Wong et al., 2018) generally operates at 60–65 °C and generates complex concatemeric products, making integration with unmodified mammalian DNMT1 potentially more challenging. Extending methylation-preserving amplification to other isothermal systems (Gill and Ghaemi, 2008; Wong et al., 2018; Tan et al., 2022) will therefore require consideration of not only temperature compatibility but also product architecture, accessory proteins and reaction chemistry.

Beyond selection of the amplification platform, an important consideration is balancing efficient maintenance of DNA methylation with suppression of DNMT1-mediated methylation of products derived from unmethylated templates. Although DNMT1 strongly prefers hemi-methylated substrates, limited activity on unmethylated DNA has been reported under certain *in vitro* conditions (Pradhan et al., 1999; Goyal et al., 2006; Adam et al., 2020). Future optimization should therefore maximize methylation recovery from methylated templates while minimizing false-positive methylation signals through systematic evaluation of enzyme concentration, reaction time, sequence context and CpG density. An additional factor that may influence methylation efficiency is the stability of the methyl donor S-adenosyl-L-methionine (SAM). SAM is susceptible to temperature-dependent degradation, and prolonged incubation above its preferred stability range may reduce the effective methyl-donor concentration while generating degradation products that inhibit methyltransferase activity (Matos and Wong, 1987; Desiderio et al., 2005). Consequently, future optimization may benefit from strategies such as staged SAM supplementation (New England Biolabs, n.d.), SAM regeneration systems or alternative reaction formats that better preserve methyl-donor availability during extended incubations.

Despite these promising results, several limitations should be addressed before MethylHDA can be considered a fully validated analytical assay. MSRE-qPCR provides a practical functional readout of methylation at predefined HpaII sites but does not directly determine methylation at every CpG within the amplified region or identify 5-methylcytosine at base resolution. In addition, the current workflow relies on a commercial HDA system optimized for short amplicons (70–120 bp) (New England Biolabs, n.d.) used in this study and was evaluated using synthetic templates representing only a limited number of methylation states. Consequently, the present study does not address genomic DNA complexity, fragmented clinical specimens, low methylated-allele fractions, amplification bias between methylated and unmethylated templates or strand-specific methylation fidelity. Furthermore, amplification and methylation were assessed primarily as endpoint measurements, leaving the temporal relationship between DNA synthesis and DNMT1 activity unresolved.

Building on these findings, future studies should evaluate multiple genomic loci spanning different GC contents, CpG densities and amplicon structures, establish calibration curves using defined methylation mixtures and determine analytical sensitivity at progressively lower DNA inputs. Base-resolution analyses, including bisulfite sequencing or enzymatic methylation sequencing (Liu et al., 2019; Sun et al., 2021), will be required to determine methylation fidelity at individual CpGs and confirm that initially unmethylated sites remain unmethylated. Direct comparison with phi29–DNMT1 amplification, conventional HDA and alternative isothermal amplification systems will further define the analytical niche of MethylHDA. Finally, validation using fragmented DNA and clinically relevant specimens will determine whether the short-amplicon design provides practical advantages for targeted biomarker detection.

## Conclusions

We developed MethylHDA, a one-step isothermal method that integrates helicase-dependent amplification (HDA) with DNA methyltransferase 1 (DNMT1)-mediated maintenance of DNA methylation in a single reaction. By establishing reaction conditions compatible with both HDA and DNMT1 activity, we demonstrated the feasibility of coupling targeted DNA amplification with concurrent maintenance of DNA methylation, as assessed by a methylation-sensitive restriction enzyme quantitative PCR (MSRE-qPCR) readout. MethylHDA establishes proof of concept for a targeted workflow for locus-specific methylation analysis and complements existing sequencing-based whole-genome methylome analysis approaches. With further optimization and validation using base-resolution methylation analyses, MethylHDA has the potential to provide a practical platform for targeted epigenetic analysis and may be extended to other isothermal amplification systems.

## Materials and Methods

### Assay to evaluate DNMT1 activity across different temperatures

To assess whether DNMT1 (Active Motif, Cat. No. 31800) retains enzymatic activity at temperatures above its optimal 37 °C, we developed an assay using a hemi-methylated DNA substrate that reports methylation-dependent cleavage by the restriction enzyme MspJI (New England Biolabs, Cat. No. R0661). DNMT1 specifically recognizes methylated cytosines on hemi-methylated DNA and catalyzes the addition of a methyl group to the complementary cytosines at CpG sites. MspJI recognizes the sequence mCNNR, where ‘mC’ is a methylated cytosine, and cleaves 12–16 nucleotides downstream. The enzyme cuts directionally: cleavage occurs only when one strand has RNmCGNR and the complementary strand has YNmCGNY, allowing MspJI to distinguish strand orientation relative to the methylated cytosine. In this system, hemi-methylated DNA is resistant to MspJI cleavage due to the absence of methylation on the recognition site. However, upon DNMT1-mediated methylation of the unmethylated strand, MspJI becomes capable of recognizing and cleaving the DNA. This provides a quantitative readout of DNMT1 activity at various temperatures.

To generate a hemi-methylated substrate that is initially unrecognizable by MspJI but becomes a cleavage target after DNMT1-mediated methylation of the reverse strand, the DNA template adapted from Petell et al. (2017) was modified as follows: the forward strand was altered at position 51 (G→T) while retaining the methylated cytosine, and the reverse strand carried a C→A mutation at position 55, with its corresponding methylated cytosine removed. These modifications create a mCGNY site on the reverse strand that is not recognized by MspJI prior to methylation. Both the DNA templates and primers were synthesized by Integrated DNA Technologies (IDT), Singapore (Table 1).

A master mix was prepared containing 1 mg/mL BSA (Sigma-Aldrich, Cat. No. A7906), 160 µM S-adenosylmethionine (SAM) (New England Biolabs, Cat. No. B9003S), 1× DNMT1 buffer (50 mM Tris-HCl, pH 8.0; 1 mM EDTA; 1 mM DTT; 5% glycerol), and 1 µM of the modified hemi-methylated DNA substrate. The mix was aliquoted into three sets of duplicate reactions. For each set, one reaction was supplemented with 425 nM recombinant DNMT1, while the corresponding control received an equal volume of water. Samples were incubated at 40 °C, 42 °C, and 45 °C for 1 hour to allow methylation to occur.

Following incubation, 1 µL from each methylation reaction was added to a digestion mixture containing either 1 unit of MspJI or no enzyme (serving as a negative control). The digestion was carried out in rCutSmart™ buffer supplemented with 0.5 µM Enzyme Activator Solution at 37 °C for 2 hours, followed by enzyme inactivation at 65 °C for 20 minutes. The resulting samples were diluted 10,000-fold and subjected to qPCR analysis.

A qPCR master mix was prepared using Maxima SYBR Green/ROX qPCR Master Mix (Thermo Fisher Scientific, Cat. No. K0221) and 0.3 µM of each primer. For each reaction, 1 µL of the diluted digestion product was added to the wells. qPCR was performed using the following thermal cycling conditions: an initial denaturation at 95 °C for 10 minutes, followed by 40 cycles of 95 °C for 15 seconds and 60 °C for 1 minute. A melt curve analysis was performed at the end of the run to verify amplification specificity and confirm the presence or absence of digestion products. Methylation efficiency was quantified by calculating the ΔCt values between digested and undigested samples.

### Assay to assess HDA activity under DNMT1-compatible conditions

To evaluate whether HDA functions efficiently under DNMT1-compatible conditions at temperatures below its optimal 65 °C, we performed an assay using the IsoAmp® II Universal tHDA Kit (New England Biolabs, Cat. No. H0110S). Enzymes and dNTP solution from the kit were incorporated into reactions formulated with DNMT1 buffer, and amplification was tested across a temperature range of 42 °C to 65 °C. Results were compared to those obtained under the standard IsoAmp® reaction conditions to assess amplification efficiency under suboptimal conditions.

Two reaction master mixes were prepared for side-by-side comparison. The first followed the standard IsoAmp® protocol with 75 nM forward and reverse primers, and 10 nM of a modified double-methylated DNA template derived from the MethylMiner™ Methylated DNA Enrichment Kit (Invitrogen, Cat. No. ME10025) (Table 2). Primers targeting the full template region were adapted from the MethylMiner™ kit. This mix was divided into two tubes: one containing IsoAmp® Enzyme Mix and the other containing water as a no-enzyme control. The second master mix was formulated using a DNMT1-compatible buffer composed of 1 mg/mL BSA, 160 µM SAM, 1× DNMT1 buffer, 4 mM MgSO₄, 40 mM NaCl, IsoAmp® dNTP solution, 75 nM primers, IsoAmp® Enzyme Mix, and 10 nM of the same DNA template. This mixture was aliquoted into three separate tubes for incubation at various test temperatures.

To assess amplification efficiency across conditions, the two reactions from the standard IsoAmp® protocol group and one reaction from the DNMT1-compatible buffer group were incubated at 65 °C for 1.5 hours. The remaining two DNMT1-buffer reactions were incubated at 42 °C, and 45 °C, respectively, for 1.5 hours. All amplification products were subsequently diluted 10,000-fold for qPCR analysis.

For qPCR, a master mix was prepared containing Maxima SYBR Green/ROX qPCR Master Mix and 0.3 µM of each primer. 1 µL of each diluted amplification product were used per reaction. The qPCR was performed under the following thermal cycling conditions: initial denaturation at 95 °C for 10 minutes, followed by 40 cycles of 95 °C for 15 seconds and 60 °C for 1 minute. Melt curve analysis was performed to confirm amplification specificity. Amplification efficiency was determined by comparing Ct values between samples with and without the IsoAmp® Enzyme Mix.

### One-pot DNA amplification and methylation assay using IsoAmp® Enzyme Mix and DNMT1

To establish a one-pot assay capable of simultaneous DNA amplification and methylation, reactions were carried out at 42 °C under DNMT1-compatible conditions. DNA templates with varying proportions of methylated and non-methylated sequences were subjected to amplification in the presence of both IsoAmp® Enzyme Mix and recombinant DNMT1, IsoAmp® Enzyme Mix alone, or neither enzyme. Following amplification, samples were treated with a methylation-sensitive restriction enzyme (MSRE) to assess whether DNMT1 had successfully introduced methylation. If methylation was preserved during the one-pot reaction, the digestion profile would reflect the methylation status of the input templates.

Methylated and non-methylated DNA templates were generated from methylated control sequences provided in the MethylMiner™ Methylated DNA Enrichment Kit (Table 2). The methylated template was modified to retain only three MSRE sites, including one HpaII recognition site (CCGG), while five methylated cytosines lacking MspJI/HpaII recognition motifs were excluded. The non-methylated template shared the same nucleotide sequence but without methylation. Primers spanning the full template region were adapted from the MethylMiner™ kit.

A reaction master mix was prepared containing 1 mg/mL BSA, 160 µM SAM, 1× DNMT1 buffer, 250 nM forward and reverse primers, 4 mM MgSO₄, and IsoAmp® dNTP solution (see Supplementary Table S1). The mixture was divided into five reactions, each containing 10 nM total DNA template with varying ratios of double-methylated and non-methylated DNA. Reactions were supplemented with 85 nM recombinant DNMT1 and/or IsoAmp® Enzyme Mix as appropriate. The five conditions were as follows: R1: 100% double-methylated template + IsoAmp® Enzyme Mix + DNMT1; R2: 50% double-methylated + 50% non-methylated template + IsoAmp® Enzyme Mix + DNMT1; R3: 100% double-methylated template + IsoAmp® Enzyme Mix (no DNMT1); R4: 100% non-methylated template (no enzyme; negative control); R5: 100% double-methylated template (no enzyme; positive control) (see Supplementary Table S2).

Reactions were incubated at 42 °C for 1 hour to enable simultaneous DNA amplification and DNMT1-mediated methylation, followed by an additional 15-minute incubation at 37 °C to facilitate completion of the methylation process. Samples were then purified using the DNA Clean & Concentrator-5 Kit (Zymo Research, Cat. No. D4014) according to the manufacturer’s protocol. Purified products were diluted 10,000-fold and used for qPCR analysis of amplification efficiency. A qPCR master mix containing Maxima SYBR Green/ROX qPCR Master Mix and 0.3 µM of each primer was prepared, and 2 µL of each diluted sample were used as input. qPCR was performed with the following thermal profile: 95 °C for 10 minutes, followed by 40 cycles of 95 °C for 15 seconds and 60 °C for 1 minute. Melt curve analysis was conducted to confirm amplification specificity.

Amplification efficiency was quantified by calculating the ΔCt between enzyme-containing and no-enzyme control reactions according to the following formula:

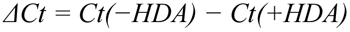

- *Ct(−HDA)* = Ct value of the no-enzyme control (without IsoAmp® Enzyme Mix)
- *Ct(+HDA)* = Ct value of the amplified sample (with IsoAmp® Enzyme Mix)

Interpretation: A large ΔCt indicates efficient amplification, as the enzyme-containing reaction produced more template (lower Ct). A small or near-zero ΔCt indicates poor or no amplification, as both reactions yielded similar Ct values.

To evaluate methylation efficiency, parallel digestion reactions were prepared by combining 5 µL of each purified sample with 5 µL of digestion buffer containing either no enzyme (undigested control) or 3 U of HpaII (New England Biolabs, Cat. No. R0171) in rCutSmart™ buffer. Reactions were incubated at 37 °C for 2 hours and then heat-inactivated at 80 °C for 20 minutes. Digested samples were diluted 100-fold for qPCR analysis using the same master mix and cycling conditions described above, with 8 µL of each diluted digestion sample added per reaction. Melt curve analysis was repeated to confirm the specificity of amplification.

Methylation efficiency was quantified by calculating the ΔCt between HpaII-digested and undigested samples according to the following formula:

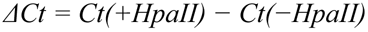

- *Ct(+HpaII)* = Ct value of the digested sample (treated with HpaII)
- *Ct(−HpaII)* = Ct value of the undigested control (no enzyme)

Interpretation: ΔCt ≈ 0 indicates complete or near-complete methylation, where the DNA is protected from HpaII digestion and amplification remains comparable to the undigested control. A positive ΔCt (> 0) indicates partial or no methylation, as HpaII digestion cleaved unmethylated DNA, reducing the amplifiable template and increasing Ct.

## Declarations

### Ethics Approval and Consent to Participate

Not applicable.

### Consent for Publication

Not applicable.

### Availability of Data and Materials

All data generated or analysed during this study are included in this published article and its supplementary information files. The datasets generated and/or analysed during the current study are available in the NCBI GenBank repository under accession numbers PZ577457 and PZ577458. The design and methylation status of the DNA templates are fully described in the Materials and Methods section.

### Author Contributions

K.W.K. contributed to brainstorming, designed and performed the experiments, analyzed the data, and wrote the manuscript. S.E.P. contributed to brainstorming, data analysis, and manuscript review. F.T.W. contributed to brainstorming and manuscript review. Y.S. contributed to brainstorming, funding acquisition, and manuscript review. W.L.C.K. supervised the study, conceived the study, contributed to experimental design and data analysis, and assisted with manuscript writing and revision. All authors read and approved the final manuscript.

## Acknowledgements

The authors thank Yi Jing Chua (Genome Institute of Singapore, Agency for Science, Technology and Research) for contributions to early-stage discussions and exploratory work.

