## Supplementary Information for "MethylHDA: One-pot helicase-dependent amplification with concurrent DNMT1-mediated maintenance of DNA methylation"

### Supplementary Figures

Figure S1

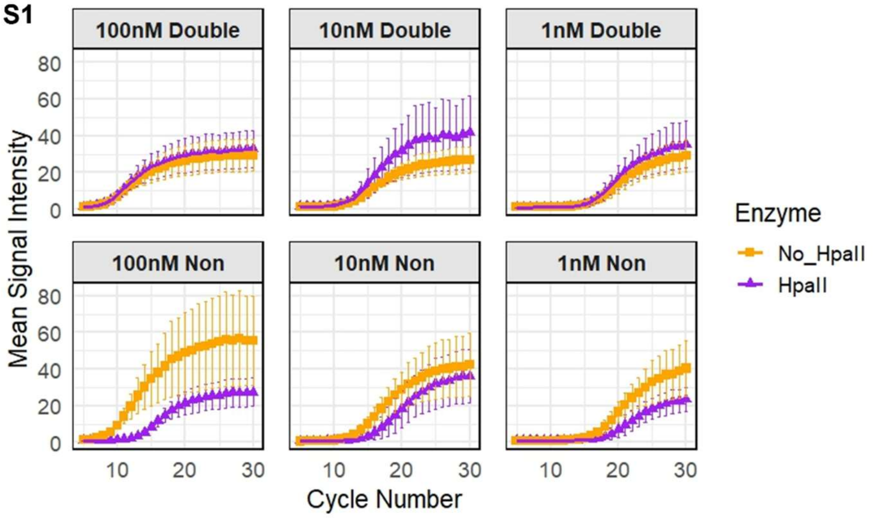


**Figure S1.** Validation of HpaII digestion efficiency across methylated and non-methylated DNA templates at different concentrations. qPCR amplification plots before and after HpaII digestion are shown for six DNA templates: 100 nM double-methylated, 100 nM non-methylated, 10 nM double-methylated, 10 nM non-methylated, 1 nM double-methylated, and 1 nM non-methylated. Templates containing non-methylated DNA show a clear shift in Ct values after HpaII digestion, indicating efficient cleavage. In contrast, double-methylated templates remain largely unchanged, confirming protection from digestion by HpaII.

Figure S2

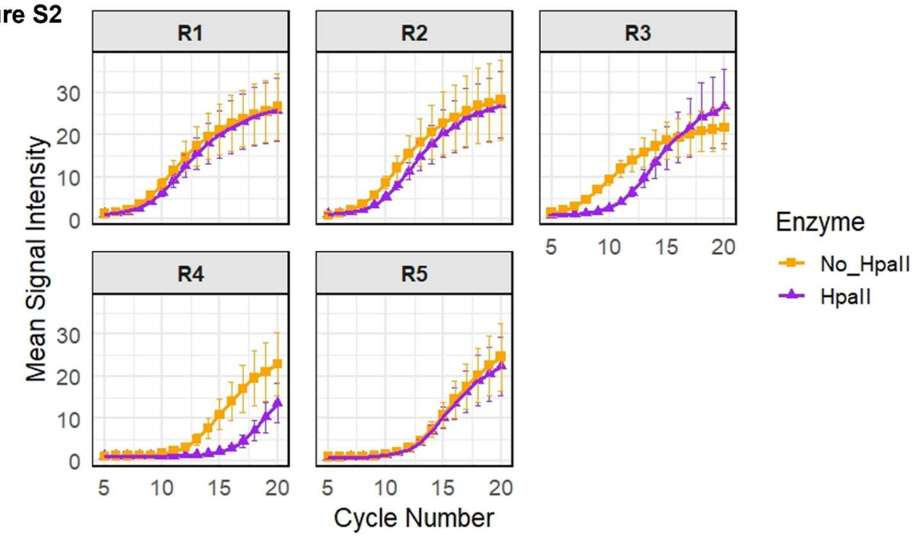

**Figure S2.** qPCR amplification curves pre- and post-HpaII digestion for samples R1–R5. DNMT1-methylated samples (R1, R2) retain strong amplification post-digestion, similar to the methylated control (R5), indicating protection from HpaII. In contrast, the unmethylated sample (R3) shows reduced amplification, comparable to the digestion-sensitive control (R4).

### Supplementary Tables

**Table S1.** Components of one-step amplification and methylation.

| Components | Amount in 20 µL reaction | Final concentration |
|---|---|---|
| 10× DNMT1 buffer | 2 µL | 1× DNMT1 buffer |
| 100 mM $MgSO_4$ | 0.8 µL | 4 mM $MgSO_4$ |
| IsoAmp® dNTP Solution | 1.4 µL | – |
| Forward and Reverse Primers (5 µM) | 1 µL | 250 nM Forward and Reverse Primers |
| 10× Bovine Serum Albumin (BSA) | 2 µL | 1× BSA |
| 1.6 mM S-Adenosyl methionine (SAM) | 2 µL | 160 µM SAM |
| IsoAmp® Enzyme Mix | 1.4 µL | – |
| 850 nM recombinant DNMT1 protein | 2 µL | 85 nM recombinant DNMT1 protein |
| 100 nM template | 2 µL | 10 nM template |
| Water | 5.4 µL | – |

**Table S2.** Composition of reactions used to assess one-pot methylated DNA amplification.

| Reaction | Templates | IsoAmp® Enzyme Mix | Recombinant DNMT1 protein |
|---|---|---|---|
| R1 | 100% Double-methylated + 0% Non-methylated | Yes | Yes |
| R2 | 50% Double-methylated + 50% Non-methylated | Yes | Yes |
| R3 | 100% Double-methylated + 0% Non-methylated | Yes | No |
| R4 | 0% Double-methylated + 100% Non-methylated (Negative control) | No | No |
| R5 | 100% Double-methylated + 0% Non-methylated (Positive control) | No | No |
